# How effective are frogs in regulating crop pest population in a natural multi-trophic system??

**DOI:** 10.1101/2021.05.26.445791

**Authors:** Deyatima Ghosh, Sabyasachi Chatterjee, Parthiba Basu

**Affiliations:** Department of Zoology, University of Calcutta, 35, Ballygunge Circular Road, Kolkata, 700-019; Ongil, 79D3,Sivaya Nagar, Reddiyar, Alagapuram, Salem, 636-004, India

**Keywords:** Frogs, Biological Pest regulation, Functional response, Intraguild predation, Trophic cascade effect

## Abstract

Potential of frogs as important natural pest control agents has been highlighted earlier. But the effectiveness of frogs in regulating the pest load in intensive agricultural landscape in a multi-trophic system is not clear. We performed controlled field experiment in paddy field with a varying density (observed in high and low agricultural intensity (AI) areas) of a commonly found frog species and compared the pest and pest predator build-up. The consumption rate of the model amphibian was studied using enclosure experiment. The consequent trophic cascade effect of frogs on both crop pest and other arthropod pest predator was analyzed using mathematical population growth models. Although frogs consumed pests, they could not reduce crop pest abundance. although a lesser frog density found in high AI areas significantly affected the pest predator abundance. Based on the functional response result, mathematical growth models demonstrated that with a constant harvesting factor (Holling’s Type II) frogs will always have a negative impact on the beneficial natural enemy population due to intraguild predation thereby limiting its potential as a pest regulator. Our study challenges the notion of frogs as an effective pest control agent and argues that increasing habitat diversity might improve overall biological pest suppression.

## 1. INTRODUCTION

Extensive use of agro-chemicals and habitat loss associated with agricultural intensification has been identified as a major driver of biodiversity loss in agricultural landscape (Dudley and Alexander, 2017; Balmford et al., 2012; Newbold et al., 2015; IUCN, 2014; Lajmanovich et al., 2003). The negative impact of biodiversity loss on Ecosystem Service (ES) delivery in agriculture has also been recognized widely (Brook et al., 2008; Tscharntke et al., 2005). Ecological Intensification (EI) of agriculture on the other hand has been suggested as a sustainable solution to conventional high external agro-chemical input-driven intensive agriculture (Tittonell, 2014). EI involves restoration or improvement of the ES provisions in agriculture e.g., crop pollination, biological control of crop pests, or soil biological activities. However, there exists a serious gap in our scientific understanding about the underlying ecological processes that generate ES (Firbank et al., 2013; Birkhofer et al., 2015; Naeem et al., 2015; Bengtsson, 2015).

Biological control of crop pests is a key ES (Bengtsson, 2015; Ives et al., 2000; Wilby and Thomas, 2002; Gurr et al., 2003). However, despite years of research, our understanding of the underlying ecological processes that govern biological control, particularly, the complex food web interactions that regulate the process, is still incomplete (Thies et al., 2005; Bengtsson, 2015). Contrary to the benefits obtained from natural enemy population, several studies have highlighted the detrimental effects it may have on the non-target population (Clarke et al., 1984; Howarth, 1983, 1991). Hence any biological pest control strategy warrants thorough scrutiny of the trophic cascade in a multi-species system.

Amphibians have long been recognized as potential natural predators for crop pests (Hamer et al., 2004; Knutson et al., 2004; Gibbs et al., 2005; Loman and Lardner, 2006) being both generalist and opportunist predator (Kathiwada, 2016; Schaefer et al., 2006; Mahan and Johnson, 2007). The importance of amphibians in biological control, particularly in rice cultivation has been particularly recognized (Li et al., 2008; Zhang et al., 2010; An et al., 2012; Zhang, 2013). A sizeable abundance of amphibians is suggested to be efficient in bringing down the rice pest population (Teng, et al, 2016; Fang et al., 2019). Kathiwada (2016) reported a high proportion of rice pests in frogs’ diet and recommended the introduction of frogs as biological pest control in rice fields. Intensive agriculture has been found to negatively impact frog population (Arntzen et al. 2017; Davies et al. 2018). Ghosh and Basu (2020) reported reduced density of amphibian fauna in areas of high agricultural intensification compared to low agricultural intensity areas. A decline in frog population in an intensive agricultural landscape is therefore expected to impact ES delivery in rice fields.

However, most studies that reported the potential of frogs as pest regulators have not looked at the effect frogs may have on non-target species especially beneficial natural arthropod pest predators. These studies also have not investigated if frog predation of rice pests could effectively bring down pest population through controlled multi-trophic field experiments.

The extent to which predation can regulate the prey population is influenced by the functional response of the predator to changing prey density (Leeuwen et al., 2007; Williams and Martinez, 2004; Wollrab and Diehl, 2015). It is therefore important to gain insights into the functional response of frogs to changing density of rice pests before coming to a conclusion about their effectiveness as rice pest regulator. However, none of the studies that prescribed frogs as potential regulators of crop pests has considered this.

The present study attempts to bridge these gaps by testing the efficiency of frogs in regulating crop pest population through a multi-trophic controlled field experiment conducted at the Agricultural Experimental Farm, University of Calcutta, West Bengal, India using a focal frog species available commonly in the agricultural fields across India. The experiment was designed with field-realistic densities of frogs found in the high and low agricultural intensity areas in the study region reported earlier by Ghosh and Basu (2020). Therefore, apart from answering if frogs are efficient regulators of rice pests, the study also explores the impact of agricultural intensification on ES delivery by frogs. We also assess the functional response of the focal frog species to varying densities of a common rice pest in the study region. We also predict if the focal frog species can be an efficient pest regulator in a multi-trophic system using a mathematical model based on our results.

## 2. Materials and Methods

### 2.1 Experiment Site

The experiment was performed in the Agricultural Experimental Farm, University of Calcutta, Baruipur Campus in West Bengal in 2018 (22.3787°N, 88.4361°E). The farm is spread across an area of 210 acres with paddy being the major crop grown during the monsoon. Total experimental area was 7,500 square meter. The follow-up mesocosm experiment to study the nature of feeding response in amphibians was performed in the University campus.

### 2.2 Experimental design

Three blocks of 50 × 50 meters area were selected to carry out the study. Each of the blocks was divided into 6 experimental plots of 10 × 10 meters where we installed our experimental units using drift fence of length 10m and height 3ft. Thus we installed a total of 18 experimental plots across an area of 7500 m^2^. We left a buffer of 5m between each experimental plot. We randomly assorted the plots to three different frog tretaments which were - one control without frogs, one with a treatment density of 10 frogs / 100 m^2^ as observed in low agricultural intensification and third with a treatment of 5 frogs / 100 m^2^ representing the density observed in high agricultural intensification zones (density data obtained from a study conducted in Odisha along an agricultural intensification gradient, Ghosh and Basu, 2020).

### 2.3 Experimental model

We used adult (5-6 cm) paddy field frogs or *Fejervarya limnocharis* as our model organism as they are the most dominant in paddy fields of India (Dash and Mahanta, 1993). They are prevalent in South Asia (Sumida et al., 2007). This species is also a generalist and feeds mainly on insects (Chuang and Borzee, 2019) hence makes it a potential bioresource to test their pest controlling efficiency. Apart from these, the species is categorized as least concern by IUCN (IUCN, Conservation International and NatureServe, 2006), hence our study did not put any undue extinction pressure. We collected amphibians by active search in August 2018 from within the campus area for release in experimental plots.

### 2.4 Experimental plot preparation

We selected paddy fields for its economic demand and the huge swathe of land under paddy cultivation and used a local rice cultivar named “Patnai”.

We prepared the seedbed on 30^th^ June at a density of 1916 seeds/ m^2^ for a total seedbed area of 210 m^2^. Paddy saplings were transplanted around 30 days after preparing seedbed from 27 to 29 July 2018 at an average density of 762 plants/100 m^2^. Before releasing the frogs, all pots were covered with a mesh net of 48mm gap size to exclude predation risk on both the pests and experimental frogs by birds as well as to prevent the escape of any frogs from the experimental enclosures. Plots were left undisturbed for 15 days before the release of frogs. Frogs were released on August 15^th^, 2018, after sunset when the temperature was low to prevent desiccation. After leaving the system undisturbed in order to allow the animals to overcome any stress during their release process, we started sampling from August 20^th^, 2018. No pesticides, herbicides, or any fertilizers were applied in the plots to eliminate any detrimental effects on animals. The study continued throughout the vegetative phase until the flower stage and crops were harvested at end of November 2018.

### 2.5 Arthropod sampling

All active and passive sampling was conducted from 6.00 hr to 13.00 hr of a day. For insect sampling, we used the sweep netting method (Fatahuddin et al., 2020) with a standard-sized sweep net of 24 inches length and an opening diameter of 12 inches. Sweep netting was done 10 times along each of the four sides of the experimental plots making a total of 40 sweeps per plot. All specimens collected were wet preserved in 70% alcohol for later identification and categorization.

#### Passive sampling: Pest Infestation study

We used a 1 × 1 meter wooden frame and placed it randomly within an experimental plot. For each sampling, data on the total number of plants and total number of leaves were collected. We selected 3 random plants within each such quadrat and inspected proportion of infestation (Teng et al., 2016) by rice hispa (*Dicladispa sp*.), leaf folder (*Cnaphalocrocis sp.*), defoliation, tungro, rice whorl maggot (*Hydrellia* sp.). For leaves where we could not identify the infestation, were collected and preserved for later identification under expert supervision. At each quadrat, we also searched for the presence of any pest or predator and wet preserved the specimen in 70% alcohol for later identification. This method was replicated 5 times for each plot per sampling.

We maintained the same sequence of sampling for the different treatments (6 plots for each treatment) and pooled observations from 6 plots per treatment in each complete sampling session. We averaged data for further statistical analysis and data representation. All the samplings were repeated four times for each plot throughout the paddy growing season. This covered all the successive stages of rice cultivation following transplantation to flowering stage.

### 2.6 Estimating the frog survival

From 17.00 hr to 21.00 hr every day we counted the frogs and checked for any dead or desiccated frog inside the experimental plot to ensure that the number of frogs remained constant.

### 2.7 Arthropod Identification

We identified all arthropods obtained in active sampling up to genus level and broadly grouped them into orders to categorize them as pests and non-pest natural predators (Khan and Pathak, 1994; NICRA, 2011) for the purpose of this present study.

### 2.8 Experiment to study the feeding potential in this species

We performed a mesocosm experiment with one size class of frogs (5-6cm, adult) and three prey densities of 8, 33, and 43. We used grasshoppers in this experiment as they are one of the major paddy defoliators. Our field observation also showed the model amphibian feeding on the grasshopper. The densities of grasshopper used were replicates from control treatments of the field experiment (observed grasshopper densities were 8, 33, 43 and 44). The entire experiment was replicated four times through the months of September till November 2019. On the day of the experiment, one predator was transferred to the experimental mesocosm of 1×1 meter that was designed to mimic the natural environment of these predators. Health of the mesocosm was maintained regularly. Total experimental time was set for 17 hr. At the end of the experiment, we removed the predator and counted the remaining prey. All predators had the same starvation period before they performed in an experiment. Since they feed on live prey we also maintained a separate terrarium for the prey at varying densities to check the mortality rate in 24 hr and found no death over the period.

To collect data on the handling time and attack rate of frogs, we replicated the experiment with varying prey densities and recorded the feeding rates for 30min with a video camera (Nikon 5200D) fitted outside the terrarium.

### 2.9 Statistical analysis

We performed a Kruskal-Wallis test to compare the difference in the number of transplanted seedlings.

Proportion of pest infestation by 5 major paddy pests was checked using nested ANOVA with a mixed-effects model where sampling phases and frog treatments were the fixed effects and treatment within blocks was included as random effects. Comparison of means was performed for both the fixed effects. We checked the significance of the random effects by comparing the above described model with a second model containing only the fixed effects. For total pest and pest predator count, a Generalized Linear Mixed Model (GLMM) with negative binomial error distribution was applied to check their abundance with varying phases of the paddy growth and frog treatments (Alexander et al., 2000; Linden and Mantyniemi, 2011; Ghosh and Basu, 2020). The model structure was the same as that of the nested ANOVA.

We performed a diagnostic test for the functional response (Pritchard et al., 2017) with prey eaten as a function of prey provided. Based on the result we created a final model with attack rate and handling time.

We used R software (version 3.5.2) (R Core Team, 2018) to perform all the analyses using packages-nlme, Mass, lmtest, multcomp, lme4, frair.

### 2.10 Mathematical Growth Model

In the presence of *n* number of frogs, the pest and pest predator dynamics will vary depending on the consumption rate of these frogs (provided the resource available to the crop pests is constant across the three treatments) and also by their mutual birth and death processes in which the predator-prey interaction plays an important role.

We designate, *x_1_* = number of pests and *x_2_* = number of pest predators while “*n*” number of frogs act as predators for both the crop pests and the crop pest predators.

Since this frog species is a generalist, it does not have any preference for crop pests or predators, and its feeding process is assumed to be a random one, with a total mass “*m*” being consumed in *“T”* time, from a “meal” that consists of prey and predator, for which the model is assumed to be as follows.

#### Model of frog predation on crop pest and predator

It is clear from the above that if frogs were to feed exclusively on pests, it would need to eat *n_1_* = *(m/m_1_)* pests while if it were to feed exclusively on the predator, it would consume *n_2_* = *(m/m_2_)* predators where *m_1_* and *m_2_* are the mass of pest and pest predator respectively. These can be satisfied with “success rates” that are proportional to *N_1_* = *(x_1_/n_1_)* and *N_2_* = *(x_2_/n_2_)* respectively and would occur with relative chances *P_1_* and *P_2_* given by,

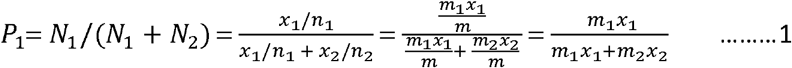

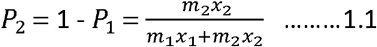

Thus with n number of frogs present, the consumption rate of pests is :

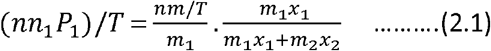

and the consumption rate for predators is

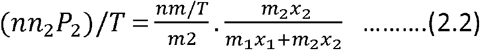

The evolution equations of the pest and predator are then,

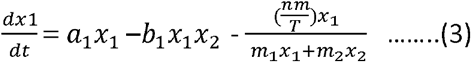

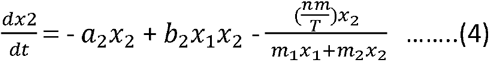

For Eq (3), the first term is the natural birth rate of the pest while the second term is their death rate due to feeding by the predator and the third term that follows from Eq (3) is the death rate of the pest due to consumption by frogs. Similarly, for Eq(4), the first term is the death rate of a predator in absence of prey while the second term is the predator’s growth rate as they feed on the prey, and the last term is the death rate of the predator due to consumption by frogs.

## 3. Results

### 3.1 Statistical Analysis

#### Control experiment

The mixed effect models showed no significance in frog treatments for any of the five major paddy pest infestation types. However their abundance varied with the growth phases of paddy crop. Table 1 provides fixed effects in models for these 5 pest infestation types. Pest abundance similarly showed a significant difference in the 2^nd^ (p < 0.000), 3^rd^ (p<0.000), and 4^th^ phase (p<0.000) of sampling but none of the frog treatments had any significant effect (5 frog treatment p=.077; 10 frog treatment p=0.369).

**Table 1:** Fixed effects (Paddy growth phase and Frog treatment) in Nested ANOVA models for the five pest infestation types

| **Pest/Disease** | **Fixed Effects** | **F value** | **p value**<sup>596</sup> |
| --- | --- | --- | --- |
| **Tungro** | Paddy growth phase | 4.92 (3,20) | 0.010* <sup>597</sup> |
|  | Frog treatment | 1.09 (2,4) | 0.41 <sup>598</sup> |
| **Defoliation** | Paddy growth phase | 6.21 (3,20) | 0.003**<sup>599</sup> |
|  | Frog treatment | 1.21 (2,4) | 0.38 <sup>600</sup> |
| **Rice whorl** | Paddy growth phase | 2.08 (3,20) | 0.13 <sup>601</sup> |
|  | Frog treatment | 1.93 (2,4) | 0.25 <sup>602</sup> |
| **Leaf folder** | Paddy growth phase | 3.88 (3,22) | 0.022* <sup>603</sup> |
|  | Frog treatment | 0.73 (2,4) | 0.53 <sup>604</sup> |
| **Rice hispa** | Paddy growth phase | 70.17 (3,20) | <.0001***<sup>605</sup> |
|  | Frog treatment | 2.32 (2,4) | 0.21 <sup>606</sup> |

Pest predator abundance appeared to be significantly affected by different phases of paddy growth i,e the 2^nd^ (p< 0.000), 3rd (p< 0.000), and 4^th^ (p=0.006) sampling phases. But, the abundance of these pest predators was also significantly affected by the lower density of frogs (5 frog treatment p=0.016; 10 frog treatment p= 0.159) (Table 2).

**Table 2:** Summary of results from GLMM for pest and insect predator build-up in response to frog treatments and paddy growth phase

| Response | Treatment |  |  | Paddy growth phase |  |  |  |
| --- | --- | --- | --- | --- | --- | --- | --- |
| Response | Control | 10 frogs | 5 frogs | 1<sup>ST</sup> | 2<sup>ND</sup> | 3<sup>RD</sup> | 4<sup>TH</sup> |
| Pest abundance |  |  | 0.09 |  | 0.000 | 0.000 | 0.006 |
| Predator abundance |  |  | 0.01 |  | 0.000 | 0.000 | 0.000 |

Figure 1 shows a prominent difference in the pest predator dynamics compared to the respective abundance in the control plot. Since the resource available for the crop pests across the treatment is the same (Kruskal Wallis chi-squared = 2.9454, df = 2, p-value = 0.229), and other external factors remaining similar, such difference in pest and predator build-up can only occur due to the harvesting factor from frogs. However, this harvesting factor is proved to be limited in the Functional Response analysis presented in the following section.

**Fig. 1.**
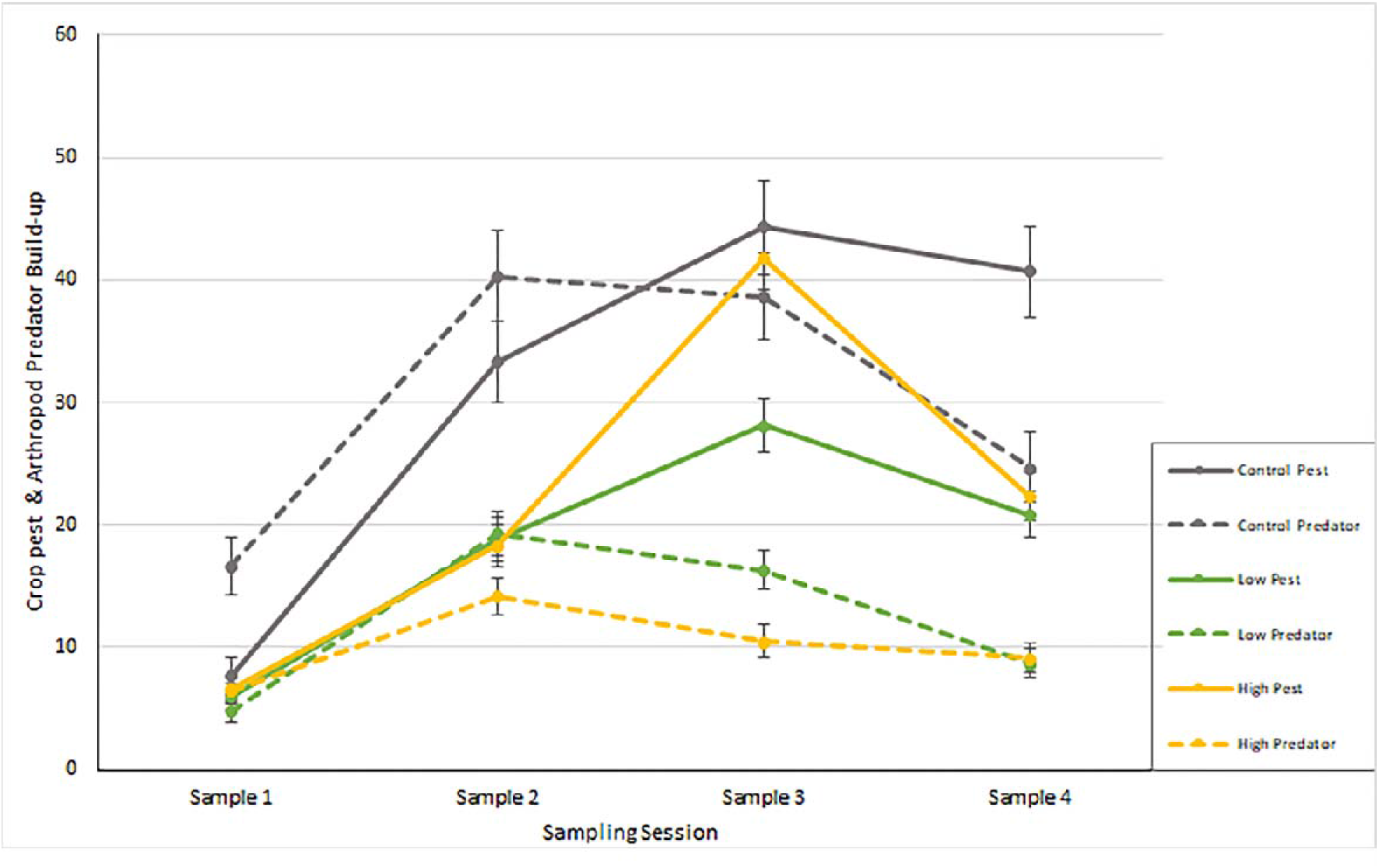
Crop pest and insect predator build-up in control, at frog densities found in High (10frogs/100 sq.m) and Low (5 frogs/100 sq.m) agricultural intensity areas at different sampling intervals.

### 3.2 Functional response from a mesocosm experiment

Diagnostic tests indicated the feeding response pattern in our focal frog species to be Holling’s Type II. We further built a model with attack rate and handling time, keeping exponential co-efficient “q” fixed at 0. The results showed a significant effect of handling time (p = 0.000) that validates our initial finding that the functional response is of Hollings’s Type II (Fig. 2). Therefore, our results show, that frogs, as predators don’t have an insatiable hunger but have a constant rate of feeding that is independent of prey density.

**Fig 2.**
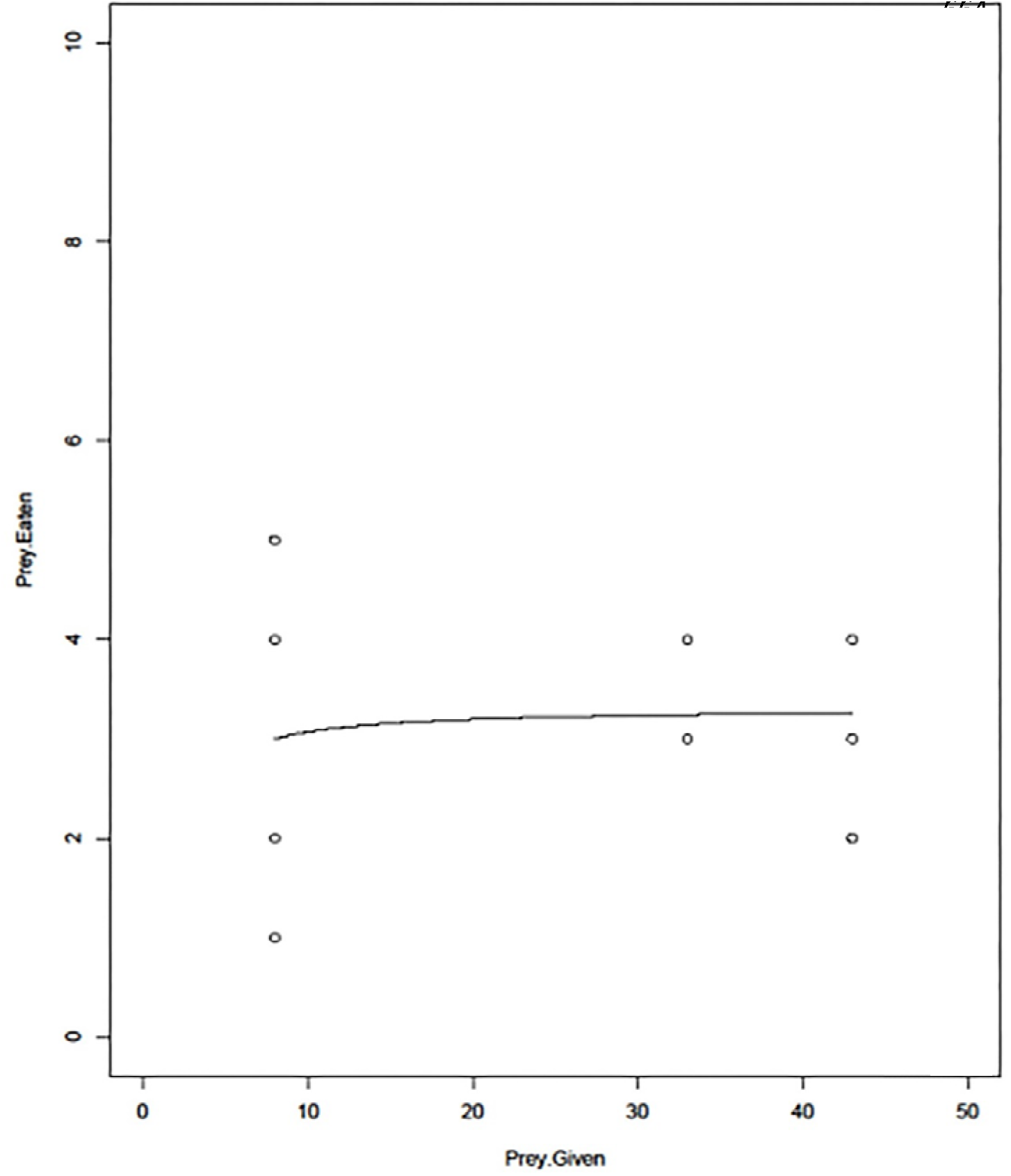
Functional response output showing Holling’s Type II feeding nature in focal frog species.

### 3.3 Assessment of trophic cascade through Mathematical growth model

For equations, 3 and 4 to be a non-trivial solution, *x*_1_≠ 0 and *x*_2_≠ 0,

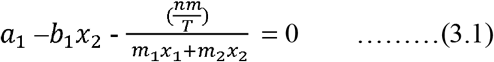

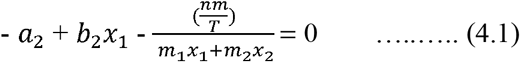

which describe the final steady-state, i.e. dx_1_/dt = 0 = dx_2_/dt. Thus on solving Eqs.(3.1,4.1) for the two unknowns, x_1_ and x_2_, we get uniquely,

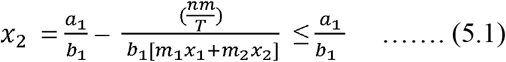

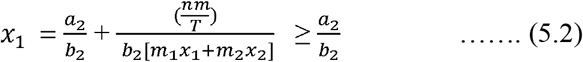

In absence of frogs, the solutions would be,

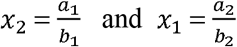

which, as expected, are also the average values 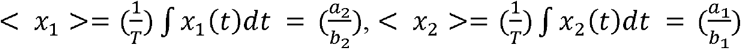 for the usual Lotka-Volterra case, around which x_1_(t) and x_2_(t) evolve in their limit cycle, where the integrals are over 0 ≤ t < T and T is sufficiently large. Our mathematical model, as of now, does not explore the possibilities of limit cycles but shows that in presence of a frog as a tertiary consumer, the trophic cascade effect would be negative for beneficial pest predators and would have positive feedback on crop pest population.

## 4. Discussion

As our study reveals, although frogs consumed pests, the existing field densities of frogs in our study region (Ghosh and Basu, 2020) are insufficient for significant pest regulation in the natural environment. This holds true for both the high and low agricultural intensification areas. Even at the highest field-realistic density of frogs as found in the low agricultural intensity areas, pest population was not affected.

That the frogs do consume insect pests have been reported earlier (Hirai and Matsui, 2002; Chen et al., 2005; Attademo et al., 2005; Yousaf et al., 2010). Teng et al. (2016) also showed that a density of 100 frogs/67sq. m and 150 frogs/ 100sq. m could effectively control pests. The ineffectiveness of frogs in controlling the insect population in our case could be due to the inadequate abundance of frogs i.e., below the threshold density for effective regulation, in the study sites.

Like any pest control strategy, the goal of a biological control strategy is to keep the pest population below an economic threshold level (ETL). Though stomach content analysis gives evidence towards amphibians contributing to pest consumption, drawing a conclusion about their ability to reduce pest load below ETL based on such information will be presumptuous. Although studies have reported a reduction in pest load in presence of frogs in a controlled setup (Teng et al., 2016; Fang et al., 2019) these studies overlooked the impact on other arthropod natural predators as found in natural settings.

We argue that frogs negatively impact the arthropod natural predators of crop pests in multiple species systems and that makes them ineffective as a biological pest regulator.

### Antagonistic interaction with other natural enemies

We observed crop pests belonging to orders Orthoptera, Coleoptera, Lepidoptera, Hymenoptera, Diptera, Hemiptera, and arthropod predators belonged to orders Odonata, Coleoptera, and Aranea. Such rice fields provide a potential ecosystem with huge prey diversity (Fernando, 1995; Guerra and Araoz, 2015).

Our study shows that, in a multiple species system amphibians exhibited a preference for pest predators rather than crop pests. Brown (1974) has reported the diet of amphibians in croplands to be dominated by non-pests (94% of diet). Similar study by Khatiwada et al. (2016) also shows amphibian diet is significantly composed of natural pest predators. Generalist predators have an antagonistic effect in biological control especially (Perez-Alvarez, Nault, & Pavedo, 2019) where the predation pressure is equal on all the prey species (Wells, 2007). The effect and strength of intraguild predation with respect to frogs have not been reported prior to this study. However, our study is the first to experimentally demonstrate the effect and strength of intraguild predation with respect to frogs.

Our mathematical growth model corroborates our result and shows that with a limited feeding rate the presence of frogs will always have a negative effect on the insect predator population when they behave as a tertiary consumer in a system with both crop pests and arthropod predators. If frogs feed on insect pests only, according to the growth equation for natural pest predators (Eq. 4) their growth rate would decrease as they are robbed off their resource, and eventually, there would be an increase in crop pest abundance. However, if the harvesting factor is shifted completely towards the crop pest predator (Eq. 3) (if they were not generalists but specialists instead, or if their feeding is influenced by the more active predators (Ahmed et al., 2016)) there still would be an increase in crop pest abundance due to release of interspecific interaction from pest predator population. Therefore the presence of frogs will cause negative feedback on the crop pest predator population consequently resulting in an increase in crop pest abundance. In intensive agricultural lands where the arthropod pest predator population is disproportionately affected (Zhao et al, 2015) or is low (Perez-Alvarez et al., 2019) amphibians could be a promising biological control agent.

Martin et al. (2013) estimated a decrease in natural pest control by 46% in landscapes that are dominated by cultivatable arable lands. In such landscapes with impaired natural pest control by insect predators, the harvesting factor of frogs will be shifted maximally to the crop pest population.

### Feeding constraints of amphibians

As our experimental result about the functional response shows, the feeding rate of *F. limnocharis* is independent of prey density and the response is significantly affected by handling time. Handling time (T_h_) is associated with every prey item consumed and is indifferent to the prey density. But even when the frog was presented with a higher density of grasshoppers, the maximum number of prey consumed was determined by the time spent in prey handling (T/T_h_). This means that the rate of prey consumption by a predator declines at higher prey densities due to handling constraints (Thorp et al., 2018). Therefore, this feeding constraint of the frog will act as a limiting factor in controlling the pest population at a higher density which will continue to grow in a single trophic interaction between crop pest and frog. For effective biological control, a considerable abundance of the amphibian species needs to be conserved in intensive agricultural lands.

Scientific management of beneficial landscape structures like semi-natural habitats, hedgerows, ephemeral water bodies, degree of connectivity with adjoining remnant natural vegetation, and habitat heterogeneity (Ghosh and Basu, 2020) enhances amphibian thriving. Complex landscapes could weaken the strength of the intraguild predatory effects by reducing niche overlap through spatial separation or even alternate resource availability (Perez-Alvarez et al., 2019). Restoring the habitat diversity can therefore be expected to promote service provisioning.

Our research significantly adds to the existing body of work exploring the potential of frogs in biological control, particularly in tropical agroecological landscapes that are heavily dependent on intensification of paddy cultivation to sustain their burgeoning populations (Aditya et al. 2020). Our study highlights the importance of studying intraguild predation effect as an important factor that determines the effectiveness of frogs as biological control agents and highlights the importance of increasing the habitat diversity in the agricultural landscape to weigh down the impacts of intraguild predation and to improve the overall ES potential of natural predators, that has implications for food and livelihood security of communities dependent on agriculture.

## Authors’ Contribution

PB and DG have conceived the idea and methodology; DG collected and analyzed data; SC and DG developed the mathematical models: PB, DG and SC contributed to writing the manuscript. All authors have made significant contributions to preparing the manuscript and give approval for publication.

## Acknowledgment

We thank Prof. Santanu Jha and his team from Bidhan Chandra Krishi Viswavidyalaya for their help in insect and pest infestation identification. We also thank Mr. Sivaraman T.N. of Ongil for his initial contribution to computational work related to the population model build-up. We acknowledge Arnab Chakraborty for his support during the fieldwork.

